# The GA4GH Variation Representation Specification (VRS): a Computational Framework for the Precise Representation and Federated Identification of Molecular Variation

**DOI:** 10.1101/2021.01.15.426843

**Authors:** Alex H Wagner, Lawrence Babb, Gil Alterovitz, Michael Baudis, Matthew Brush, Daniel L Cameron, Melissa Cline, Malachi Griffith, Obi L Griffith, Sarah Hunt, David Kreda, Jennifer Lee, Javier Lopez, Eric Moyer, Tristan Nelson, Ronak Y Patel, Kevin Riehle, Peter N Robinson, Shawn Rynearson, Helen Schuilenburg, Kirill Tsukanov, Brian Walsh, Melissa Konopko, Heidi Rehm, Andrew D Yates, Robert R Freimuth, Reece K Hart

## Abstract

Maximizing the personal, public, research, and clinical value of genomic information will require that clinicians, researchers, and testing laboratories exchange genetic variation data reliably. Developed by a partnership among national information resource providers, public initiatives, and diagnostic testing laboratories under the auspices of the Global Alliance for Genomics and Health (GA4GH), the Variation Representation Specification (VRS, pronounced “verse”) is an extensible framework for the semantically precise and computable representation of variation that complements contemporary human-readable and flat file standards for variation representation. VRS objects are designed to be semantically precise representations of variation, and leverage this design to enable unique, federated identification of molecular variation. We describe the components of this framework, including the terminology and information model, schema, data sharing conventions, and a reference implementation, each of which is intended to be broadly useful and freely available for community use. The specification, documentation, examples, and community links are available at https://vrs.ga4gh.org/.

## Introduction

Precision medicine and contemporary biomedical research are increasingly driven by large, coordinated genome-guided efforts^1–12^. The analysis of patient genomic data in the clinical setting has enabled tremendous advances in health care delivery through genome-guided diagnosis and clinical decision support^13–16^. However, numerous technical, financial, and legal obstacles impede the adoption of genomic science on a global scale. To address these challenges, the Global Alliance for Genomics and Health (GA4GH) was formed as a policy-setting and technical standards development organization, and now comprises over 600 leading organizations in the domains of healthcare, research, patient advocacy, life science, and information technology^17^. GA4GH brings together expertise from a diverse international set of real-world, genomic-data-sharing Driver Projects. These Driver Projects contribute to domain-specific teams, or Work Streams, to promote the sharing of health and genomic data according to the Findable, Accessible, Interoperable, and Reusable (FAIR) principles^18^.

Ensuring that precision genomic medicine is effective for individuals and for health systems will require that clinicians, researchers, and testing laboratories communicate genomic variation and related information reliably. Although widely-adopted standards for certain classes of variation already exist, many of these formats have been purpose-built for specific applications, including human-readable standards such as the Human Genome Variation Society (HGVS) variant nomenclature^19^, the International System of Human Cytogenomic Nomenclature (ISCN)^20^, and the PharmVar Pharmacogenetics nomenclature^21^, as well as genome-oriented flat file formats such as the Variant Call Format (VCF)^22^, among others (**Supplemental Table 1**). All current standards have design constraints that preclude a comprehensive coverage of variation types and extensibility to new types.

In response to this need, the GA4GH Genomic Knowledge Standards (GKS) Work Stream led the development of the Variation Representation Specification (VRS, pronounced “verse”), a community-driven and extensible specification to standardize the exchange of diverse variation data. Throughout the specification and this manuscript, we use the term *variation* to mean the molecular or quantitative state of a referenced biological molecule. This very broad definition is in contrast with the commonly used *variant* term, which we define as an alternate sequence state when compared to a reference sequence. This distinction sets the scope for VRS, which complements existing variant representation standards and provides an expressive mechanism for computational representation of variation. While many current standards (e.g. ISCN or HGVS nomenclatures) are designed to be amenable to visual interpretation by humans, VRS focuses on computational precision, expressiveness, and extensibility rather than human readability. As a result, VRS is more verbose than other contemporary human-readable variant representations, but better suited to expressing and representing complex variation concepts. VRS is a natural complement of human-readable nomenclatures when used for the exchange of genomic information from databases, clinical reports, or scientific manuscripts. VRS currently covers many classes of variation that are defined on a contiguous molecule such as single nucleotide variants (SNVs), multi-nucleotide variants (MNVs), indels, repeats, and haplotypes. VRS also supports variation describing absolute and relative abundance within a system, such as transcript expression and copy number variation. While VRS is intended to be extensible to virtually any form of variation, initial efforts have focused on those types of variation of greatest impact to the biomedical research and clinical genomics communities. Similarly, VRS is readily usable for description of variation in non-human systems.

As we move towards global federation of clinical genomics resources, the development of standards supporting a federated model of variation representation is increasingly important. Here we describe the components of VRS and their use in enabling a federated system of resources for the functional and clinical annotation of variation. We summarize the key components enabling the precise and extensible representation of variation with VRS, including the underlying terminology, information model, schema, and conventions for computing globally consistent identifiers.

### The Variation Representation Specification

To achieve a precise and computable representation of variation, VRS comprises several interdependent components, including a terminology and information model, machine-readable schema, sharing conventions, and globally unique computed identifiers (**Figure 1**). These components allow for the specification to address multiple use cases and to evolve to suit the needs of the community.

**Figure 1.**
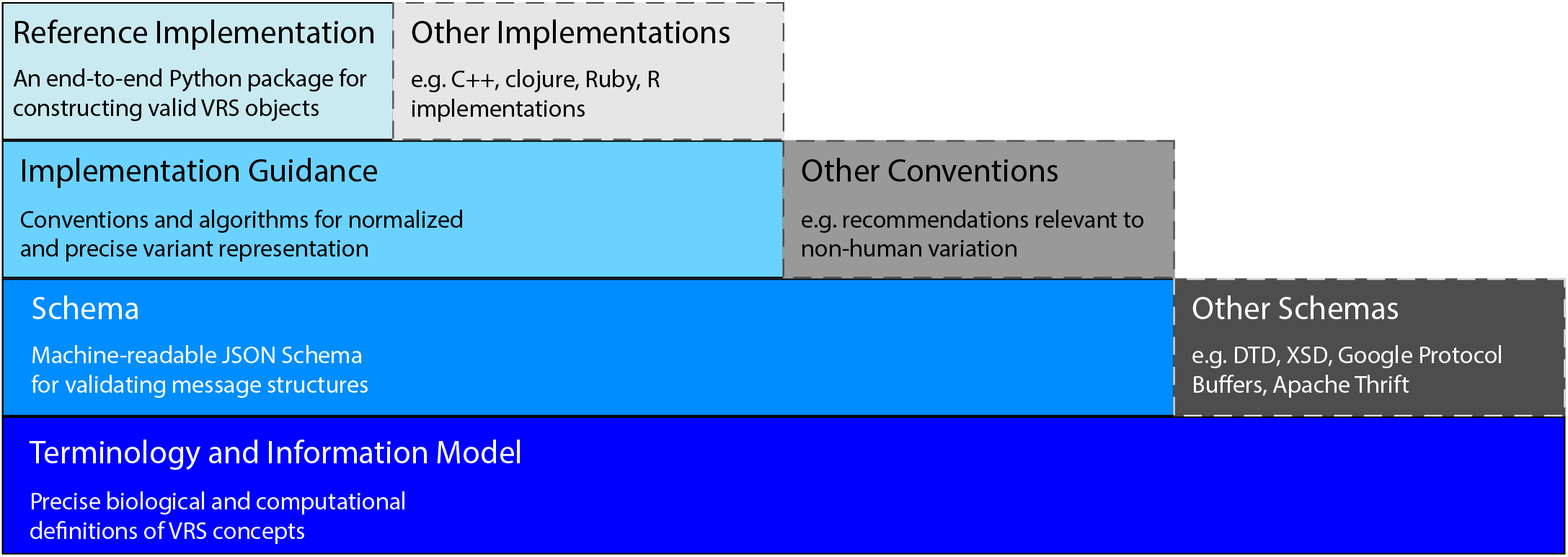
Components and Extensions of the Variation Representation Specification. VRS is a specification comprising multiple layered components (solid border blue boxes) that each serve as a foundation for the components above it. While VRS provides a full end-to-end framework, each component of the framework can be extended by the community into alternate forms (dash border gray boxes). For example, VRS provides a JSON Schema implementation based upon the Terminology and Information Model, but that same Terminology and Information Model may be used to build schemas in DTD, XSD, Google Protocol Buffers, Apache Thrift, or other data validation frameworks. This modular construction of VRS encourages interoperability across many scenarios and communities.

In totality, VRS provides a framework for the unique and semantically precise characterization of variation. To achieve this, VRS objects are *Value Objects*–minimal information objects intrinsically defined by their content (e.g. a C>T transition in residue 2 of the sequence CCTAC)–rather than records for human presentation from system identifiers (e.g. clinvar:13961). This design choice is fundamental to each component of the specification; objects are immutable and have a specific meaning prescribed by the attributes they contain. Values extraneous to the meaning of a VRS object, such as the name of a location (“chr 7”) or a label for a well-known variant (“BRAF V600E”) are not captured directly within VRS objects.

#### The scope of VRS and other GA4GH genomic knowledge standards

While VRS provides a framework for precise, computable representation of variation, there are several closely related challenges in genomic knowledge representation that are beyond the scope of this standard. The first of these challenges is the selection of a representative *variant context* that defines the subject of a biological annotation. Many public variant registries (e.g. CIViC^23^, ClinGen Allele Registry^11^, COSMIC^6^, and ClinVar^3^) aggregate multiple different variant contexts under controlled identifiers. For example, a genomic variant on multiple genome assemblies, the associated transcript changes, and predicted protein changes are each variant contexts that may all be linked under a single “variant” ID in such resources. Integrating knowledge from resources that aggregate variant contexts is challenging, and processes for selecting and/or normalizing to a particular context is a policy decision that is beyond the scope of this work. The GA4GH GKS Work Stream is developing a Variant Origination Policy to help variant registries address these challenges.

By virtue of being value objects, VRS objects do not contain human-readable labels (e.g. transcript and chromosome identifiers, HGVS variant descriptions, or concept aliases), or data that can be recreated from the provided message (e.g. reference sequence). While trivial to implement on a per-exchange basis, additional specifications for the types and values of these common data descriptors are necessary to standardize the exchange of these data between systems that share them. The GKS Work Stream is currently developing a Value Object Descriptor framework to standardize this type of information exchange.

Finally, VRS does not support the characterization or capture of attributes relevant to experimental conditions. This means that VRS does not specify how to represent characteristics of the assay (e.g. microarray, whole-genome sequencing, gel electrophoresis), measurement (e.g. read count, signal intensity), or inferred characteristics (e.g. sample ploidy, clonality, purity) beyond the variation itself. To support these cases, alongside other statements of biological and clinical evidence, the GKS Work Stream is building a larger framework to support variant annotations (https://github.com/ga4gh/va-spec).

#### Terminology and information model

The VRS terminology and information model, informed by community authorities such as the ISCN and Sequence Ontology^24^, is the foundation for the VRS Schema. Definitions of biological terms in the scientific community may be abstract or intentionally ambiguous, reflecting imprecise or uncertain measures due to limitations in our understanding of those concepts. Occasionally this creates divergent usage of terms across communities. For example, the term *genotype* has common meanings as either the alleles at a single genetic locus or as a collection of alleles at several genetic loci. While humans can readily discern between these overloaded definitions from contextual observations, abstract and ambiguous terms are not readily translatable into a computable representation of knowledge. Therefore, the specification begins with precise computational definitions for biological concepts that are essential to representing sequence variation. The VRS information model specifies how the computational definitions are represented semantically as inter-related objects and how values are to be represented in fields.

An important distinction made in VRS is between variation on a single resultant molecule (*molecular variation*) from variation that pertains to a quantitative evaluation of molecules (*systemic variation*). Molecular variation includes substitutions, insertions, deletions, haplotypes, and structural rearrangements. Systemic variation, in contrast, is used to describe gene expression variation, copy number gain/loss variation, and genotypes (planned) that may involve several molecules within a system (e.g., genome). For example, the HGVS expression “NC_000001.10:g.(?_15764950)(15765020_?)dup” has been used by some clinical labs to describe a copy number gain, assayed by a method that cannot confirm that the gain is a result of a tandem duplication. However the use of the dup syntax, by definition in the HGVS recommendations, characterizes this event as a tandem duplication. These two variation concepts — a tandem duplication on a molecule and a systemic copy number gain — are often described ambiguously, leading to potential misinterpretation of descriptions of variation by data consumers. Importantly, VRS also maintains the separation of concerns between these two statement types; if a data source intends to specify a tandem duplication *and* specify a systemic copy number gain as a result of that duplication, these are represented as two distinct variation objects associated with the corresponding sample by that source. Wide adoption of VRS will enable systems to appropriately distinguish molecular and systemic copy number changes, from which derived human-readable forms (e.g. a CNV representation or an HGVS dup) may be generated.

VRS enables genomic data providers to clearly delineate these two types of variation and several other common variation concepts. This is enabled through precise technical definitions of classes for MolecularVariation (*Allele*, *Haplotype*) and SystemicVariation (*Abundance*) concepts. Additional technical Variation concepts (*VariationSet*, *Text*) and concepts supporting these variation types (*Location*, *State*, and *Interval*) are also defined. Text variation serves as a useful mechanism for an interim representation of variants that are otherwise not yet representable with VRS. Some examples of these are covered in our planned concepts under active development, including structural variation, genotypes, and transcript locations (**Table 1**). Technical definitions for primitive concepts supporting these objects (*CURIE, Residue, Sequence*) are also provided.

**Table 1.** VRS Objects.

| Object Type | VRS Identifier Prefix | VRS Release | Purpose |
| --- | --- | --- | --- |
| **Variation** |  |  |  |
| Allele | VA | v1.0 | Contiguous insertions and deletions at a specific Location |
| Text | VT | v1.0 | Other forms of variation (technical compatibility) |
| VariationSet | VS | v1.1 | Collections of variation |
| Haplotypes | VH | v1.1 | Phased molecular variation |
| AbsoluteAbundance | VAA | v1.2 | An absolute systemic quantity / copy number variation |
| RelativeAbundance | VRA | v1.2 | A relative systemic quantity / copy number variation |
| Genotypes |  | Planned | Summary of variation corresponding to a Location |
| Structural Variation |  | Planned | Novel molecules composed from sequence originating at non-contiguous locations |
| **Location** |  |  |  |
| SequenceLocation | VSL | v1.0 | Locations on defined IUPAC character sequences |
| ChromosomeLocation | VCL | v1.1 | Locations on defined cytogenetic regions |
| **SequenceExpression** |  |  |  |
| LiteralSequence | N/A | v1.2 | Used to define a specific sequence of literal IUPAC characters |
| DerivedSequence | N/A | v1.2 | Used to define a sequence described by a sequence location |
| RepeatedSequence | N/A | v1.2 | Used to define a specific sequence by a number of repeats of an indicated subsequence |
| **Interval** |  |  |  |
| SimpleInterval | N/A | v1.0 | An interval on a sequence specified by a start and end coordinate |
| CytobandInterval | N/A | v1.1 | A cytogenetic interval specified using cytoband nomenclature |
| NestedInterval | N/A | v1.2 | A compound interval specified by a set of inner and outer coordinates |

Variation, Location, State, and Interval are all extensible abstract base classes (**Figure 2**). This feature of the specification enables extensibility of the model, such as the re-use of Location classes to describe not only sequence variation, but also variation described by a gene concept or a cytoband. Similarly, while Alleles will often be composed from a *SequenceLocation* (a location defined by an interval on a residue sequence) and a *LiteralSequence* (a character string of IUPAC character codes^25^), the newer *RepeatedSequence* enables composition of Alleles that may have clinical significance when represented as a repeating subsequence, such as measuring CAG repeats in Huntington’s Disease^26^ and microsatellite instability in colorectal cancers^27^. The flexibility of these classes allows for the semantically precise representation of many common forms of variation, currently including substitutions, insertions, deletions, phased variation, and copy number variation.

**Figure 2.**
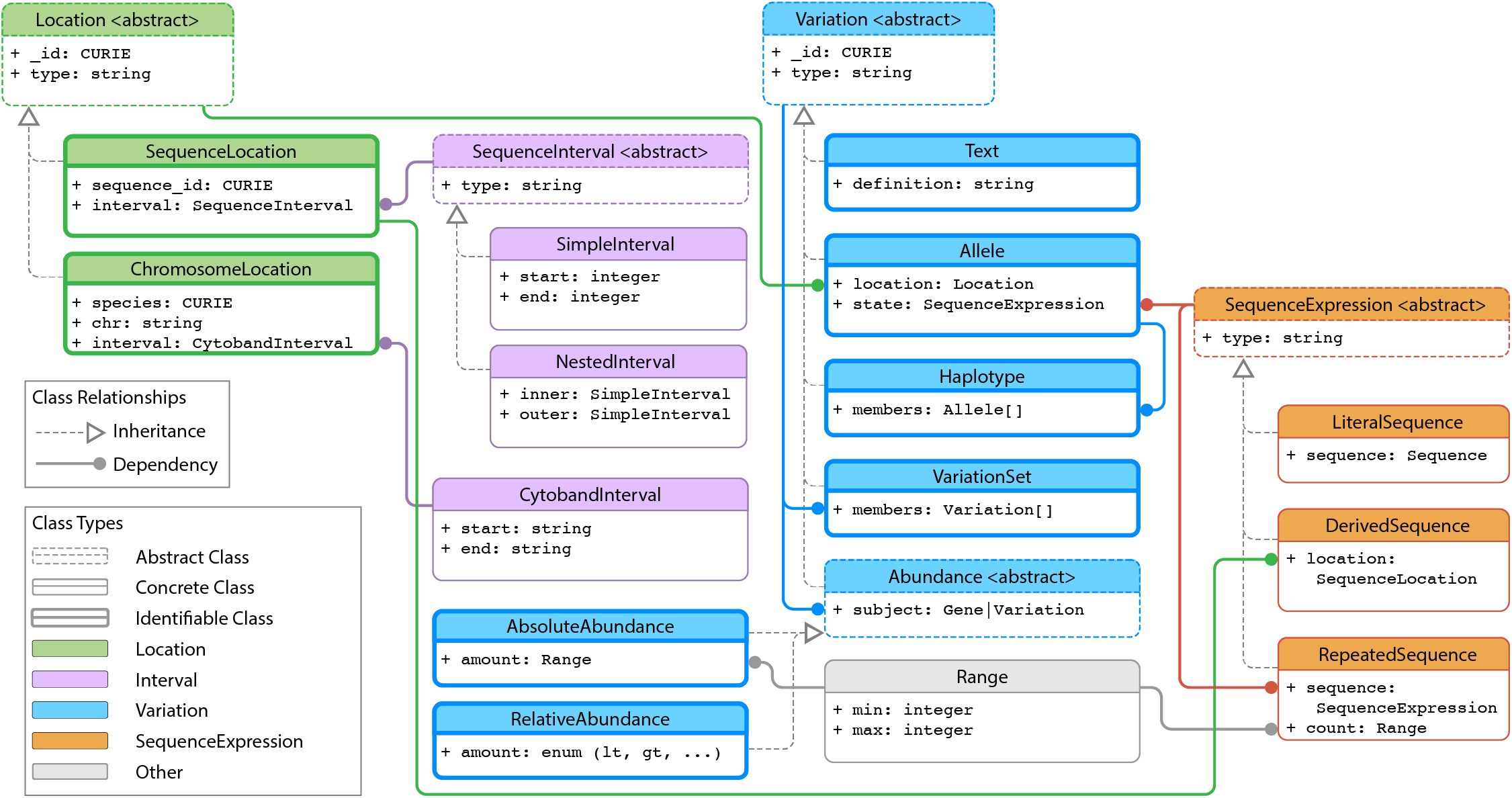
Information Model. The VRS information model consists of several interdependent data classes, including both concrete classes (solid borders) and abstract superclasses (dashed borders). These classes may be broadly categorized as conceptual representations of *Location* (green boxes), *Variation* (blue boxes), *SequenceExpression* (orange boxes), and *Interval* (purple boxes). Other objects support these primary classes, including Range (gray box), and GA4GH *Sequence* strings (not shown). While all VRS objects are Value Objects, only some objects are intended to be identifiable (bold borders). Relationships between objects are indicated by connecting lines for class inheritance (dashed line arrows) and attribute dependency (solid line arrows with circular arrowheads).

#### Machine-readable schema

To be useful for information exchange, the information model must be realized in a structured message syntax. VRS specifies its message syntax through a schema currently implemented in JSON Schema. However, the VRS information model could be readily translated to other schema frameworks (e.g. DTD, XSD, Google Protocol Buffers, Apache Thrift) as desired. Use of the schema enables data consumers to validate VRS objects that are passed between systems. The schema defines the attributes and associated value types for valid VRS objects, and it also includes regular expression validation for relevant attributes (e.g. compact URIs and cytoband descriptions). The VRS repository includes language-agnostic tests for ensuring schema compliance in downstream implementations.

#### Conventions that promote reliable data sharing

Building upon the terminology, information model, and schema, VRS also provides recommended conventions regarding the generation of VRS objects to best facilitate data sharing. While the schema provides the structure of messages, these conventions assist in the evaluation and selection of values to use in VRS objects.

A key convention of VRS is the use of inter-residue coordinates for specifying sequence locations. Inter-residue coordinate conventions are used in this terminology because they provide a level of conceptual consistency that is not possible with residue coordinate systems. While natural for humans, residue coordinates have a critical shortcoming when dealing with sequence variation. Specifically, interval coordinates are interpreted as exclusive coordinates for insertions, but as inclusive coordinates for substitutions and deletions (**Figure 3a**). As a consequence, the use of residue coordinates requires knowledge of the operation type (insertion / deletion / substitution) in order to understand the meaning of the indicated positions. By choosing an inter-residue coordinate system, VRS is able to construct Location objects that have a singular, immutable interpretation regardless of the variation context.

**Figure 3.**
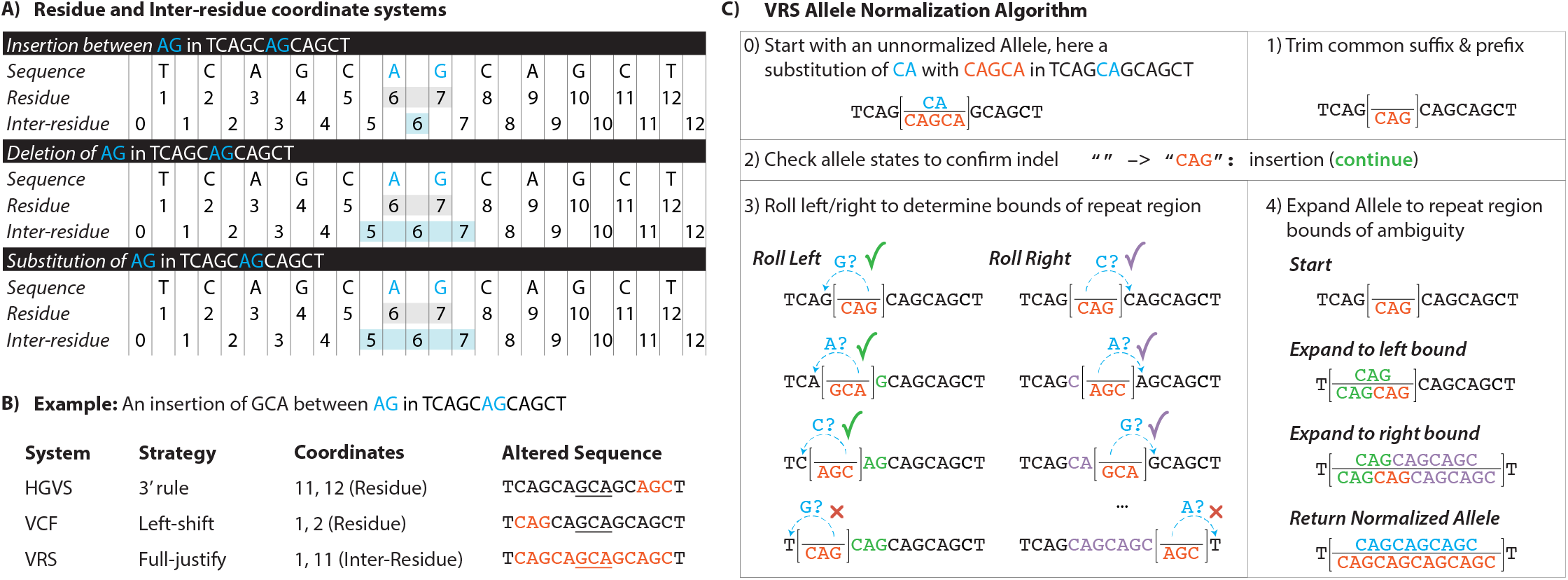
VRS Conventions. VRS provides many conventions to precisely describe and normalize molecular variation. **(A)** One key difference between VRS and existing genomic variant formats such as HGVS^19^ and VCF^22^ is the use of inter-residue coordinates. In this example, the same residue coordinates are used to ambiguously describe a two nucleotide sequence (for deletions and substitutions) and the space between those two nucleotides (for insertions). Inter-residue coordinates allow for precise representation of nucleotide or inter-nucleotide position without requiring knowledge of the operation, decoupling location representation from the representation of variation. **(B)** Here a three-nucleotide insertion occurs in a repetitive region, creating ambiguity as to where the true event (underlined) actually occurred. Three systems for describing this variant are depicted. In HGVS, the 3’-most insertion is selected as the normalized insertion producing this variant (orange lettering); in VCF, the left-most insertion is selected. In contrast, VRS avoids the selection of an arbitrary over-precise representation, and instead uses a full-justification representation that covers the entire region of ambiguity. Coordinates are the location of the insertion on the reference sequence depicted in panel A, using residue or inter-residue coordinates as specified by the corresponding representation system. **(C)** Full-justification Allele normalization is enabled by a specified normalization algorithm extended from VOCA^28^. In this example, the unnormalized Allele “reference” and “alternate” sequences (step 0) are trimmed of their common suffix “CA” (step 1). Only the resulting “reference” sequence is blank, indicating this is an insertion, and the algorithm continues (step 2). The non-blank “alternate” sequence is incrementally rolled left to identify the left bound of matching repetitive sequence, then incrementally rolled right to identify the right bound (step 3). These boundaries are used to prepend and append the regions of ambiguity to both sequences, resulting in a normalized, fully-justified Allele (step 4).

Readers familiar with these two sequence coordinate systems may also know inter-residue coordinates as “0-based”, a synonymous term used in the sequence variation community. We avoid the use of this term in our specification in favor of inter-residue, to clarify the intention of this coordinate system to members of the community new to these concepts. We have made this decision because the description of coordinates as “0-based” risks potential confusion with a hypothetical system where coordinates refer to residues, but arbitrarily begins sequences at coordinate 0 instead of the common convention of 1. We encourage community adoption of the term inter-residue to reduce these potential miscommunication errors.

Some classes of variation, such as an insertion or deletion in a repeat region, may have ambiguous representations. That is, the same empirical resulting sequence could be represented with multiple variation expressions. Normalization is the process by which a representation is converted into a single canonical form^11,28–30^. VRS adopts the use of a fully-justified representation (**Figure 3b**), ensuring that insertions and deletions in repetitive regions are not arbitrarily located to a specific position within a sequence, but instead describe the alteration over the entire region of ambiguity. This is achieved through the VRS Allele Normalization algorithm (**Figure 3c**; **Supplemental Methods**), which is an extension of NCBI’s Variant Overprecision Correction Algorithm (VOCA)^28^. Notably, the normalization method used to describe ambiguous insertions in repetitive regions only applies to Literal Sequence insertions, analogous to the use of the 3’ rule for HGVS insertions in these regions. Both VRS and HGVS also provide alternate representations to define specific repeated subsequences in these regions.

#### Globally unique computed identifiers

VRS provides an algorithmic solution to deterministically generate a globally unique identifier from a VRS object itself. All valid implementations of the VRS Computed Identifier will generate the same identifier when the objects are identical, and will generate different identifiers when they are not. The VRS computed identifier scheme works for all classes of variation in VRS and is intended to be used by other GA4GH specifications, such as the Refget API Specification (https://samtools.github.io/hts-specs/refget.html).

The algorithm for constructing VRS identifiers consists of four operations (**Figure 4a**, **Supplemental Methods**). First, a VRS object is normalized using type-specific rules defined in VRS; normalization applies, in principle, to all object types in order to standardize representation. Second, the object is serialized (i.e. converted into a string) using well-defined rules in VRS, based on a canonical JSON format. Third, a digest of the serialized data is created using the common SHA512 hashing algorithm and truncating the output to 24 bytes, which are subsequently encoded using the IETF standard base64url character set. Finally, the computed digests are prepended by a class-specific identifier prefix (**Table 1**) and the ga4gh namespace prefix. Identifier prefixes are intended to persist with the underlying data model and terminology, providing transparency into the class of object being described and analogous to the use of ENST prefixes for Ensembl/GENCODE transcript identifiers. Using a 24-byte digest virtually guarantees the uniqueness of variation identifiers; for example, in a hypothetical collection of 10^18^ objects, the probability of a single collision is less than 10^2212;21 31^.

**Figure 4.**
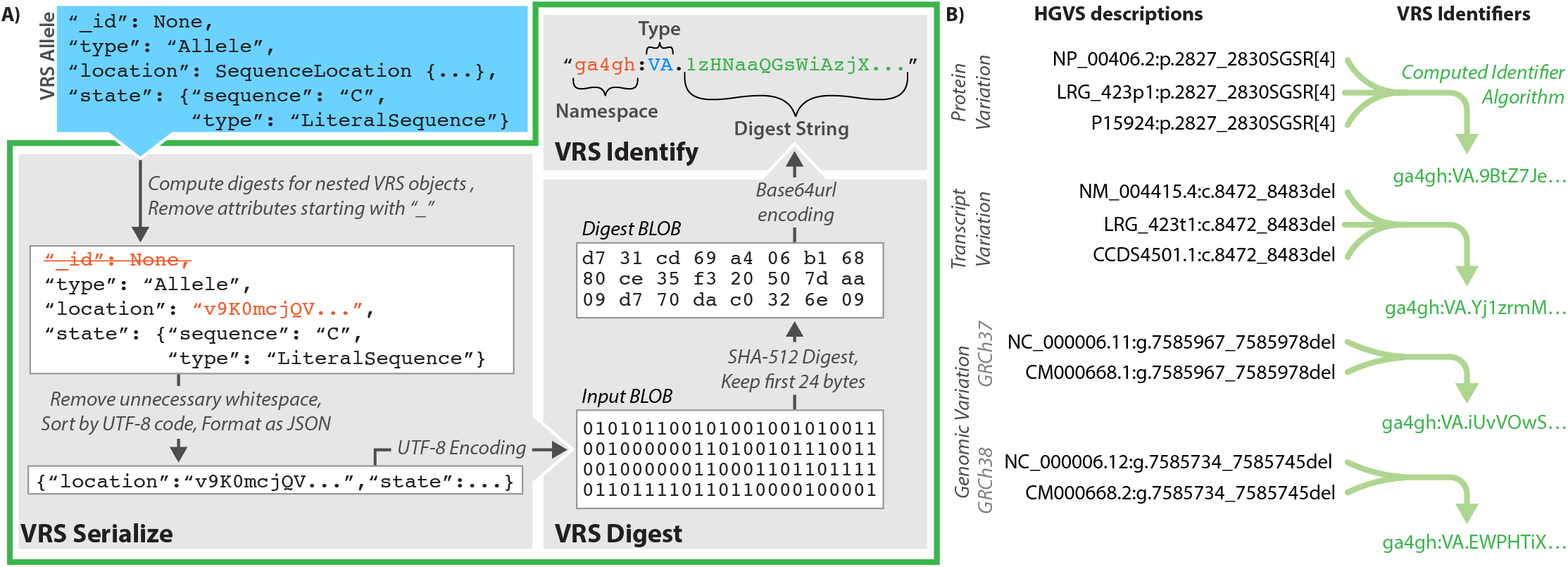
Computed identifiers. VRS provides a mechanism for federated variation identification via the Computed Identifier Algorithm. **(A)** The Computed Identifier Algorithm is defined in three stages. First, an identifiable VRS object such as an Allele (blue box) is transformed into a well-defined and canonical serialized JSON representation. The serialized Binary Large Object (BLOB) is then digested via the SHA-512 algorithm, truncated to retain only the first 24 bytes, and subsequently encoded using base64url. The resulting digest string (green text) is then appended to the object type identifier; for an Allele object, the identifier prefix is “VA” (blue text). The identifier is then assembled into a compact URI (CURIE) under the ga4gh namespace (orange text). **(B)** Use of the VRS framework enables de-duplication of identical variation concepts with differing HGVS descriptions. Here, multiple synonymous HGVS descriptions are indicated for a variant on genome builds GRCh37 and GRCh38, the corresponding transcript variant, and predicted protein translation. These four contexts (two genome assemblies, transcript, and protein) resolve to four distinct identifiers, regardless of which synonymous description is used to build the VRS object. Ellipses (“…”) used in objects and strings in this diagram represent content that is omitted for simplicity of presentation. For additional details, see https://vrs.ga4gh.org/en/stable/impl-guide/computed_identifiers.html.

VRS variation identifiers depend on underlying sequence identity rather than human assigned accessions. As a result, semantically equivalent variation objects that are defined on synonymous accessions will result in the same VRS identifier (**Figure 4b**). VRS computed identifiers are amenable to arbitrary reference sequences, including proprietary sequences or segments of a graph genome. The VRS computed identifier algorithm obviates pre-negotiation of variant identifier namespaces by allowing computational pipelines to generate reliable private identifiers efficiently and on-demand. This lowers barriers for distributed groups to share and collate interpretations with computed identifiers as keys.

#### Implementations and Community Adoption

We provide a Python package (VRS-Python) that implements the above schema and algorithms, and supports translation of VRS to and from the HGVS and SPDI variant representation schemes to facilitate rapid adoption for genomic data sharing. Support for VCF translation will be addressed in a future version of VRS-Python. VRS has been adopted by, and is being evaluated by, several major institutions in the bioinformatics community, including the ClinGen Allele Registry, the NCBI, the BRCA Exchange, and the VICC MetaKB. Both VRS (https://github.com/ga4gh/vrs) and VRS-Python (https://github.com/ga4gh/vrs-python) are publicly available on GitHub and maintained under the Apache v2.0 license. While VRS-Python may be used as the basis for development in Python, it is not required in order to use VRS. Additional community implementations of VRS have been created in C++ (**Supplemental Table 2**). Community feedback, requests, and contributions to these repositories are welcome and encouraged. Documentation is automatically generated from the VRS repository and made available online for reference at https://vrs.ga4gh.org/.

## Conclusions

The Variation Representation Specification is a GA4GH-approved standard, developed through the coordinated effort of variation representation experts and major genomic data providers from industry, government, and academic sectors. It is intended to support exchange of genomic variation data between computational systems, with a focus on semantic precision, extensibility, and conventions to promote reliable, federated identification and search.

At its foundation, VRS provides a terminology and information model. Use of these components and the decision to design VRS objects as Value Objects are a novel approach to variation representation that provides data consumers the necessary tools to reliably send and reconstruct the immutable and precise semantic meaning of a given variant. Importantly, this is done using only the minimal information content provided within VRS objects. Adherence to semantic versioning gives data consumers confidence that VRS objects will be semantically consistent and fully compatible with any future v1.x extensions of the specification. In addition, the terminology is freely available, open-source, and permissively licensed, simplifying the use and reuse of VRS objects.

Beyond the information model, VRS provides a schema and implementation guidance to promote variation messages that are consistent across implementations. A language-agnostic test suite and open-source VRS-Python reference implementation are key tools freely available to the community to reduce barriers to entry. In addition, the VRS Computed Identifier Algorithm enables a federated network for the exchange of genomic variation, through the generation of unique computed identifiers. The algorithm is enabled by the VRS information model and normalization conventions, and allows genomic data providers to consistently identify variation without prior negotiation between resources. This enables free exchange of variation data as part of a federated network and reduces the normalization burden on downstream data consumers. Together, these features of VRS provide a framework for interoperability, reinforced by the growing network of services that have implemented VRS-compliant API endpoints.

While the design choices leading to the precise, computable representation of variation through VRS enables new opportunities for genomic data sharing, these decisions also bring new considerations for genomic data providers, and leave additional challenges to be addressed. Labels that are used to describe sequences (e.g. “NC_000001.11”, “chr 1”, “NM_001374258.1”), a single variant (e.g., “SCV000504256”), or sets of variation (e.g. “VCV000013961.13”, “CA123643”, “rs113488022”, “deltaF508”) are extraneous to the minimal information used to construct VRS objects, and so resources wishing to provide descriptors for these concepts must transmit them in parallel with VRS objects or reference endpoints where these descriptions may be retrieved. In practice, GA4GH Driver Projects using VRS have found it desirable to use VRS objects to precisely represent variation, and wrap VRS objects inside *Value Object Descriptors* (VODs) to provide the various human-readable labels describing VRS objects.

A related challenge is that many genomic variation registries aggregate several related variant contexts under a single identifier. Reuse of these resource-provided identifiers then requires downstream data consumers to tease apart the intent of the identifier from the aggregate contexts, a non-trivial exercise. This challenge is compounded when trying to integrate information from multiple sources that make different choices for constructing sets of variant contexts.

The GA4GH GKS Work Stream is working to build complementary standards to VRS to address these challenges. In close collaboration with GA4GH Driver Projects, we are developing a policy for the selection and description of the originating context from aggregate variation identifiers, which is analogous to other community policies such as the *Matched Annotation from NCBI and EMBL-EBI* (MANE) Select transcript set (https://www.ncbi.nlm.nih.gov/refseq/MANE/). We are also developing a formal specification for Value Object Descriptors, in close coordination with the emerging GA4GH Variation Annotation specification. GKS is also investigating policies and frameworks by which VRS objects can be attached and referenced in written works, to enable the precision and extensibility of computational variation representation with VRS to accompany free-form descriptions in natural language.

As a specification and framework for the federated exchange of genomic variation data, the GA4GH Variation Representation Specification is precise, reproducible, and extensible to all forms of biomolecular variation. It separates concerns between genomic location and state, as well as molecular and systemic forms of variation. provides an integrative collection of components for the description, representation, and validation of variation concepts between systems. It is accompanied by multiple implementations from major genomic data providers, including an open-source and freely available reference Python implementation. The latest version of the specification is freely available for reference online at https://vrs.ga4gh.org/.

## Acknowledgements

The authors thank Christopher Bizon (Renaissance Computing Institute), Karen Eilbeck (University of Utah), Cristina Y. Gonzalez, Tim Hefferon (NCBI), Brad Holmes (NCBI), Anna Lu (National Cancer Institute), Donna R. Maglott (NCBI), Christa Lese Martin (Geisinger), and Lon Phan (NCBI) for important discussions and critical feedback that substantially advanced this work. The authors also thank Ewan Birney (GA4GH, European Molecular Biology Laboratory, European Bioinformatics Institute), Peter Goodhand (GA4GH), and Angela Page (GA4GH) for providing organizational support that enabled this work.

## Author Contributions

Author contributions follow the Contributor Roles Taxonomy (CRediT) conventions. AHW, LB, GA, MiB, MaB, DLC, MC, OLG, SH, DK, JeL, JaL, EM, TN, RYP, KR, SR, HS, KT, ADY, RRF and RKH contributed to Conceptualization. AHW, LB, GA, DLC, JeL, TN, SR, KT, RRF and RKH contributed to Methodology. AHW, LB, MC, EM, TN, RYP, KR, SR, BW and RKH contributed to Software. RYP, KR and BW contributed to Validation. AHW, LB, DLC and RKH contributed to Formal analysis. AHW, LB, GA, EM, TN, RRF and RKH contributed to Investigation. AHW, LB, GA, MiB, MC, RYP and KR contributed to Resources. AHW, LB, MiB and RKH contributed to Data Curation. AHW and RKH contributed to Writing - Original Draft. AHW, LB, GA, MiB, MaB, MC, MG, OLG, SH, DK, JeL, JaL, EM, TN, KR, PNR, HS, KT, MK, HR, ADY, RRF and RKH contributed to Writing - Review & Editing. AHW, LB, RRF and RKH contributed to Visualization. LB, MiB, MG, OLG, MK, HR, ADY and RRF contributed to Supervision. AHW, LB, MiB, MK, HR, ADY, RRF and RKH contributed to Project administration.

