## Supplemental Methods for "The GA4GH Variation Representation Specification (VRS): a Computational Framework for the Precise Representation and Federated Identification of Molecular Variation"

### VRS Allele Normalization Algorithm

The VRS Allele Normalization Algorithm is a procedure by which all Alleles–reference-matched, substitutions, insertions, and deletions–are normalized to a precise, unambiguous form. This algorithm was designed for Allele instances in which the *Reference Allele Sequence* and *Alternate Allele Sequence* are precisely known and intended to be normalized. In some instances, this may not be desired, e.g. faithfully maintaining a sequence represented as a repeating subsequence through a *RepeatSequence* object. We also anticipate that these edge cases will not be common, and encourage adopters to use the VRS Allele Normalization Algorithm whenever possible.

The VRS Normalization Algorithm is defined as follows:

0. Start with an unnormalized Allele, with corresponding “reference” and "alternate" Allele Sequences.
   a. The *Reference Allele Sequence* refers to the subsequence at the Allele SequenceLocation.
   b. The *Alternate Allele Sequence* refers to the Sequence described by the Allele state attribute.
   c. Let *start* and *end* initially be the *start* and *end* of the Allele SequenceLocation.
1. Trim common flanking sequence from Allele sequences.
   a. Trim common suffix sequence (if any) from both of the Allele Sequences and decrement *end* by the length of the trimmed suffix.
   b. Trim common prefix sequence (if any) from both of the Allele Sequences and increment *start* by the length of the trimmed prefix.
2. Compare the two Allele sequences, if:
   a. both are empty, the input Allele is a reference Allele. Return the input Allele unmodified.
   b. both are non-empty, the input Allele has been normalized to a substitution. Return a new Allele with the modified *start*, *end*, and *Alternate Allele Sequence*.
   c. one is empty, the input Allele is an insertion (empty *reference sequence*) or a deletion (empty *alternate sequence*). Continue to step 3.
3. Determine bounds of ambiguity.
   a. Left roll: Set a *left_roll_bound* equal to *start*. While the terminal base of the non-empty Allele sequence is equal to the base preceding the *left_roll_bound*, decrement *left_roll_bound* and circularly permute the Allele sequence by removing the last character of the Allele sequence, then prepending the character to the resulting Allele sequence.
   b. Right roll: Set a *right_roll_bound* equal to *start*. While the terminal base of the non-empty Allele sequence is equal to the base following the *right_roll_bound*, increment *right_roll_bound* and circularly permute the Allele sequence by removing the first character of the Allele sequence, then appending the character to the resulting Allele sequence.
4. Construct a new Allele covering the entire region of ambiguity.
   a. Prepend characters from *left_roll_bound* to *start* to both Allele Sequences.
   b. Append characters from *start* to *right_roll_bound* to both Allele Sequences.
   c. Set *start* to *left_roll_bound* and *end* to *right_roll_bound*, and return a new Allele with the modified *start*, *end*, and *Alternate Allele Sequence*.

### VRS Serialization Procedure

Digest serialization converts a VRS object into a binary representation in preparation for computing a digest of the object. The Digest Serialization specification ensures that all implementations serialize variation objects identically, and therefore that the digests will also be identical. VRS provides validation tests to ensure compliance.

Although several proposals exist for serializing arbitrary data in a consistent manner, none have been ratified. The VRS Serialization Procedure for creating computed identifiers is distinct from JSON serialization or other serialization forms. Although Digest Serialization and JSON serialization appear similar, they are NOT interchangeable and will generate different GA4GH Digests. As a result, VRS defines a custom serialization format that is consistent with these proposals but does not rely on them for definition; it is hoped that a future ratified standard will be forward compatible with the process described here.

The first step in serialization is to generate message content. If the object is a string representing a Sequence, the serialization is the UTF-8 encoding of the string. Because this is a common operation, implementations are strongly encouraged to precompute GA4GH sequence identifiers as described in Required External Data.

- If the object is a composite VRS object, implementations MUST:
- ensure that objects are referenced with identifiers in the ga4gh namespace
- replace nested identifiable objects (i.e., objects that have id properties) with their corresponding digests
- order arrays of digests and ids by Unicode Character Set values
- filter out fields that start with underscore (e.g., _id)
- filter out fields with null values

The second step is to JSON serialize the message content with the following REQUIRED constraints:

- encode the serialization in UTF-8
- exclude insignificant whitespace, as defined in RFC8259§2
- order all keys by Unicode Character Set values
- use two-char escape codes when available, as defined in RFC8259§7

The criteria for the digest serialization method was that it must be relatively easy and reliable to implement in any common computer language.
